# Effect of cyclic loading on the ultimate tensile strength of small metallic suture anchors used for attaching artificial tendons in rabbits

**DOI:** 10.1101/2024.06.08.597378

**Authors:** Obinna P. Fidelis, Pierre-Yves Mulon, David E. Anderson, Dustin L. Crouch

## Abstract

**Background:** Suture anchor failures can lead to revision surgeries which are costly and burdensome for patients. The durability of musculoskeletal reconstructions is therefore partly affected by the design of the suture anchors.

**Purpose:** The purpose of the study was to quantify the strength of different suture anchors whose sizes are suitable for attaching artificial Achilles and tibialis cranialis tendons in a rabbit model, as well as determine the effect of cyclic loading on the anchoring strength.

**Method:** Four anchors (two with embedded eyelet and two with raised eyelet, n=5 per group) were tested with cyclical loading (1000 cycles and 4.5 mm/sec) and without cycling, to inform the failure loads and mode of failure of the suture anchors. An eyebolt screw with smooth eyelet was used as a control for the test groups.

**Results:** All samples in all groups completed 1000 cycles and failed via suture breakage in both test conditions. All anchors had failure loads exceeding the peak Achilles tendon force in rabbits during hopping gait. The data analysis showed an effect of anchor type on the maximum tensile force at failure (*F*_*max*_) in all suture categories but not an effect of loading condition. Also, the Anika anchor had a significantly less adverse effect on suture strength compared to Arthrex anchor (p=0.015), IMEX anchor (p=0.004) and Jorvet anchor (p<0.001). We observed a greater percentage of failure at the mid-section for the anchors with the raised eyelets compared to the anchors with embedded eyelets, which all failed at the knot.

**Conclusion:** Anchors with embedded eyelets had clinically preferred mode of failure with less adverse effects on suture and, may be more reliable than anchors with raised eyelets for attaching artificial Achilles and tibialis cranialis tendons in rabbits.

## INTRODUCTION

Clinically, suture anchors are used to reconnect soft tissues, such as tendons and ligaments, to bone. They can provide fast, dependable, and stable anchoring of the soft tissue, facilitating healing and reunion to the bone (Burnham et al., 2020; Cho et al., 2021; Ravin et al., 2005). The strength of suture anchors is superior to alternative methods for fixing soft tissues to bone, such as bone tunnels, staples, screws, washers, and tapered plugs (Barber et al., 1993). Suture anchors have different designs and use different materials including metals and polymers. They also can be classified as screw-type and non-screw-type (Barber et al., 1995). The screw-type anchors often have larger load-to-failure limits than their non-screw-type equivalents (Barber et al., 2006; Barber et al., 2003). Metallic screw-type suture anchors are frequently used because they are biocompatible, relatively inexpensive, and provide long-term stabilization as the tissue heals (Longo et al., 2019).

Suture anchor failure, either via anchor pullout or suture breakage, is a clinically important concern because failure can result in devasting complications and pose a challenge to the success of orthopaedic surgeries (Ntalos et al., 2021). In a recent study (Fidelis et al., 2024), we used a veterinary metallic screw-type anchor (2.7 mm x 9 mm, Part No. 60-27-09, IMEX Veterinary Inc., Longview, TX) with either USP size 1, 2 or 5 braided composite sutures (Fiberwire, Arthrex, Inc., Naples, FL) to attach artificial Achilles tendons to the calcaneus bone in rabbits. In nine of twelve rabbits, the suture failed; the timing of failure were determined to occur within the first 10 weeks after implantation based on radiographic observation and confirmed by post-mortem dissection. In eight of nine specimens with failure, the suture broke at the mid-section, away from the knot. The knot is considered the weakest part of tied sutures and the failure mode distant from the knot suggested that the failure was caused by the suture either wearing or cutting on the edge of the anchor eyelet. This hypothesis is further supported by the fact that the reported average peak Achilles tendon force in laboratory rabbits during hopping (57.7 N, West et al., 2004) is much lower than the reported knot strength (295 N) of the size 5 suture (Arthrex Inc, 2014). Based on our studies and the available literature, further engineering analysis is needed to evaluate risk factors for failure modes.

Several factors, acting either independently or interactively, may have caused the observed suture failures. Two of these factors can be generally categorized as (1) eyelet design and (2) loading conditions. The eyelet is the hole in the anchor through which the suture passes. In most metallic screw-type anchors, the eyelet is either raised above or embedded within the screw (Table 1). The geometry of the eyelet, including the radius of edges, also can vary among anchors. These eyelet features can affect the relative motion between the suture and anchor as well as the concentration and magnitude of reaction forces (e.g., normal, shear, friction) on the suture. The IMEX veterinary anchor we used in our previous study to attach the artificial Achilles tendon had a raised eyelet, and some edges on the raised post of the anchor near the eyelet have relatively acute, “sharp”, angles (Figure 1). Sharp edges reportedly reduce the failure loads of sutures around the eyelet by up to 73 percent compared to suture contact with a smooth surface (Meyer et al., 2002; Wasik et al., 2013).

**Table 1.**
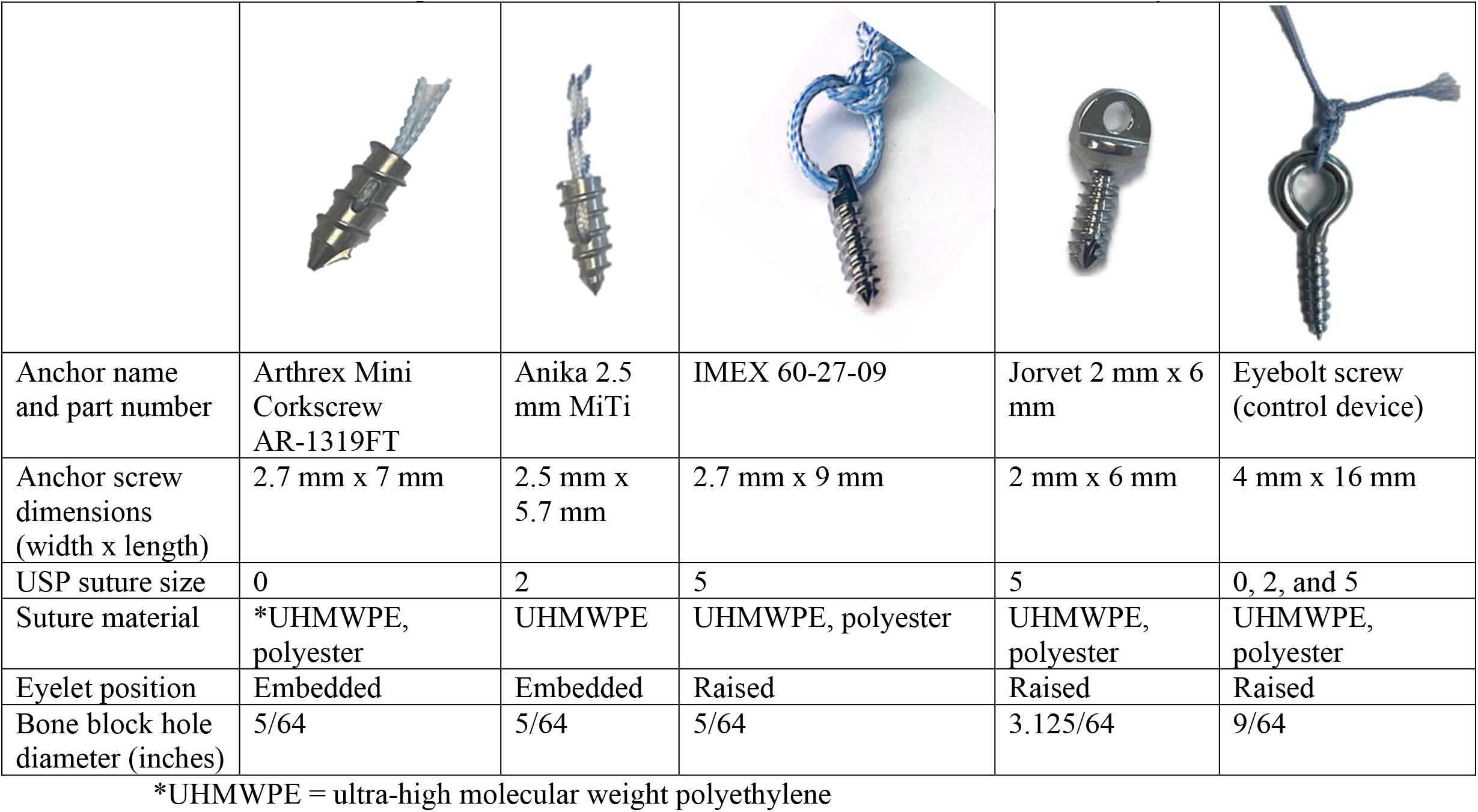
specifications and features of suture anchors tested in this study.

|  |  |  |  |  |  |
| --- | --- | --- | --- | --- | --- |
| Anchor name and part number | Arthrex Mini Corkscrew AR-1319FT | Anika 2.5 mm MiTi | IMEX 60-27-09 | Jorvet 2 mm x 6 mm | Eyebolt screw (control device) |
| Anchor screw dimensions (width x length) | 2.7 mm x 7 mm | 2.5 mm x 5.7 mm | 2.7 mm x 9 mm | 2 mm x 6 mm | 4 mm x 16 mm |
| USP suture size | 0 | 2 | 5 | 5 | 0, 2, and 5 |
| Suture material | *UHMWPE, polyester | UHMWPE | UHMWPE, polyester | UHMWPE, polyester | UHMWPE, polyester |
| Eyelet position | Embedded | Embedded | Raised | Raised | Raised |
| Bone block hole diameter (inches) | 5/64 | 5/64 | 5/64 | 3.125/64 | 9/64 |

\*UHMWPE = ultra-high molecular weight polyethylene

**Figure 1.**
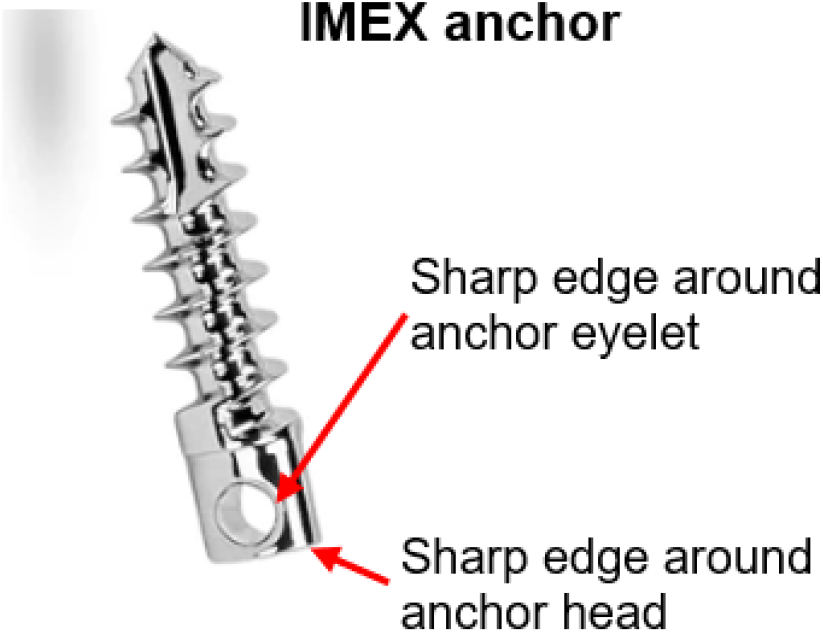
IMEX anchor highlighting the sharp edges around the anchor’s eyelet and head.

Previous studies investigated the effects of eyelet design and loading conditions on suture anchor strength, but not for select anchors whose sizes we have identified as being compatible with rabbit models used for attaching artificial Achilles tendons to the calcaneus. Since design characteristics and mechanical properties can vary widely among suture anchors, results from a subset of suture anchors may not be able to be generalized to all. Therefore, the objective of our study was to determine the effect of eyelet design and cyclic loading on suture strength for four metallic screw-type suture anchors, two with raised eyelets and two with embedded eyelets, whose sizes are compatible with the rabbit model. An eyebolt with a smooth cylindrical cross-section was used as a control. We hypothesized that 1) the suture anchors with a raised eyelet would have an adverse effect on maximum tensile force at failure (*F*_*max*_) when compared with suture anchors having an embedded eyelet; and 2) cyclical loading would have a more pronounced adverse effect on *F*_*max*_ for the suture anchors with a raised eyelet than for the suture anchors with an embedded eyelet; 3) a greater percentage of failure mode would occur at the suture mid-section (i.e., distant from the knot) in the suture anchors with a raised eyelet than in the suture anchors with an embedded eyelet.

## METHODS

### Suture Anchors

We tested four screw-type, metallic suture anchors that are of suitable size for attaching artificial Achilles tendons to the calcaneus in rabbits (Table 1). Two anchor types had raised eyelets and two anchor types had embedded eyelets. The two anchors with raised eyelets, both marketed for veterinary use only and made of stainless steel, were used in our previous in vivo rabbit study: one (2.7 mm x 9 mm, Part No. 60-27-09, IMEX Inc., Longview, TX) to attach an Achilles artificial tendon to the calcaneus bone and the other (“Jorvet” 2 mm x 6mm suture screw, cortical, SKU J0836D, Jorgensen Laboratories LLC., Loveland, CO) to attach a tibialis cranialis artificial tendon to the 5^th^ metatarsal bone. The two anchors with embedded eyelets were made of titanium and marketed for use in humans. Since we suspected that the edges of the raised eyelets may contribute to suture wear during cyclic loading, we used an eyebolt screw (4 mm x 16 mm, Home Depot, Atlanta, GA) whose eyelet had a smooth, circular cross-section (Table 1) as a control group; the radius of curvature of the eyebolt eyelet was ∼3.375mm.

In all anchors, we used the largest suture size that the anchor could accommodate. The Arthrex Mini Corkscrew anchor is typically packaged with a USP size 2-0 braided composite suture, but we manually changed the suture to the largest size that the anchor could accommodate (size 0). For the Anika anchor, we used the suture that was supplied with the anchor (Parcus Braid #2 suture, Anika, Bedford, MA); the suture is composed of ultra-high molecular weight polyethylene (UHMWPE). In all other anchors, we used a #5 braided composite suture composed of a UHMWPE core with a braided polyester covering (Fiberwire, Arthrex, Inc., Naples, FL). All 3 suture sizes were tested using the control eyebolt screw.

### Study Design

Each of the four anchors and eyebolt screw listed in Table 1 were subjected to two different loading conditions (n=5 samples per condition): (1) cyclic loading followed by load to failure and (2) single cycle loading to failure. For the Jorvet anchor, five different samples were tested in both loading conditions. For the remaining anchors, ten different anchors were used so that each anchor was tested only once. The same five eyebolt screws were used to test both loading conditions for all three suture sizes. The order of anchor or loading condition was not randomized.

### Mechanical Testing Procedure

Each anchor was screwed into pre-drilled holes (hole diameter listed in Table 1) on a simulated bone block (PCF 50, density=0.8 g/cm^3^, Sawbones, Vashon, WA); the holes were drilled perpendicular to the block surface. We used one hole per anchor per test to avoid poor block-anchor grip. Each anchor was accompanied with a braided composite suture according to the size listed in Table 1. The suture was tied around a braided polyester rope that represented the artificial tendon but whose diameter (6.5 mm) was substantially larger than that of the artificial tendon to ensure that the suture would fail first, as observed in vivo. The suture was tied with six throws of square knots (Tidwell et al., 2012). All samples were clamped and tested on a universal material testing machine (Model 5965, Instron Inc., Norwood, MA), that has a load cell with static rating ± 5kN and deflection at force capacity of 0.12 mm.

All loading was applied in line with the central long axis of the suture anchor (180°) as shown in Figure 2. The suture anchors were preloaded to 60 N, the approximate peak Achilles tendon tensile force in rabbits during hopping gait (West et al., 2004). The samples were then unloaded and preloaded to approximately 1 N prior to testing. For single cycle to failure loading tests, the samples were loaded to failure under continuously increasing force until failure. For samples subjected to the cyclical loading condition, samples were loaded uniaxially for 1000 cycles from 0 N to 60 N, then ramped up to failure. The extension rate for all tests, 4.5 mm/s, was approximately the greatest extension rate that the testing machine could apply and still track the target cyclic load limits accurately. This extension rate was chosen to mimic the rapid extension/loading rate expected in the rabbit hindlimb Achilles tendon during hopping gait.

**Figure 2.**
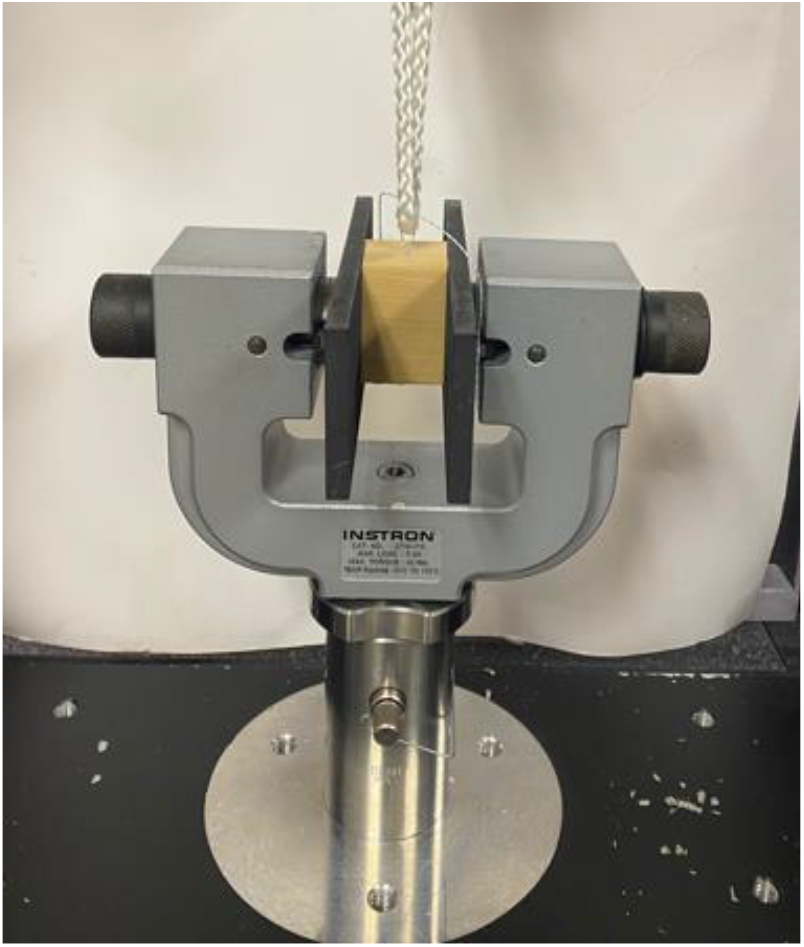
Prepared test specimen clamped in the universal material testing machine.

### Statistical Analysis

From the time-series force data for each test, we extracted the maximum tensile force at failure (*F*_*max*_). A test of the normality of the data for each category of suture was performed using the standardized residuals of the failure loads and a Kolmogorov-Smirnov test. An initial two-way analysis of variance (ANOVA) was performed for each suture size category (size 0, 2, and 5) with factors ‘anchor type’ and ‘loading condition’ followed by Tukey HSD pair-wise comparison between each anchor and eyebolt of similar suture size.

To permit direct comparison among suture anchors with different suture sizes, for each loading condition, we calculated the percent difference in *F*_*max*_ for each suture anchor test sample (*F*_*max*,*anchor*_) with respect to the mean *F*_*max*_ of the eyebolt screw 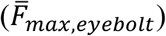 with the corresponding suture size:

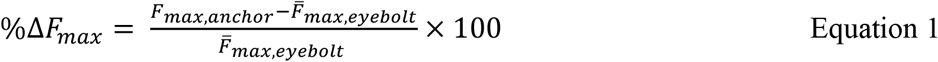

%Δ*F*_*max*_ indicates the extent to which the suture anchor affects *F*_*max*_ relative to the “ideal” eyebolt screw. Using the data calculated from equation 1, we performed a second two-way ANOVA test across all anchors with factors “anchor type” and “loading condition”, followed by Tukey HSD post-hoc pairwise comparisons. We used IBM SPSS statistics (v.28) to perform the normality test and ANOVA. Fisher’s exact test was used to compare the proportion of suture failure modes (suture breakage at “knot” vs “mid-section”) among groups. For all statistical comparisons, p<0.05 was considered significant.

## RESULTS

All samples completed 1000 cycles without failure. Furthermore, all suture anchors failed by breakage of the suture rather than anchor pull-out from the bone block. In this way, the failure mode (suture breakage) was consistent with that observed in our previous in vivo study. The results show that for all samples in both loading conditions, the tensile force (load) increased to the maximum and then rapidly dropped after suture breakage (Figure 3).

**Figure 3.**
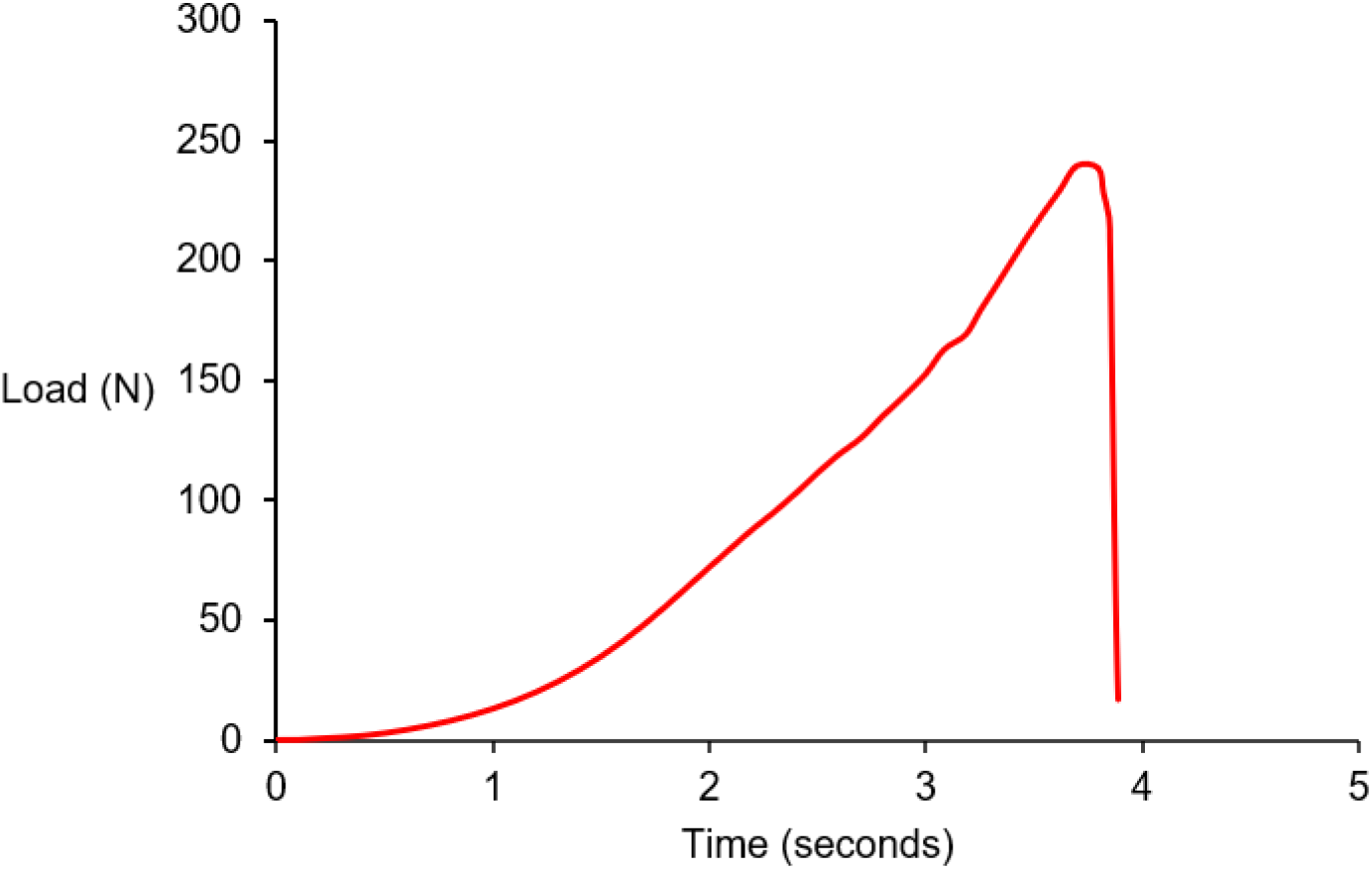
Load-time curve of a sample of the Anika anchor loaded to failure without cycling. The maximum tensile force at failure for this sample equals 239.51 N.

The results show that data for *F*_*max*_ for each suture category was not significantly different (suture size 0, p=0.084; suture size 2, p=0.200; sutures size 5, p=0.200) from a normal distribution. For the % reduction in suture strength using the *F*_*max*_ of the eyebolt corresponding to each anchor, the data also was not significantly different from a normal distribution for Arthrex (p=0.138), Anika (p=0.200), IMEX (0.190) and Jorvet (p=0.200).

*F*_*max*_ for all anchors and loading conditions exceeded the reported peak Achille tendon force in rabbits during hopping gait (57.7 N, West et al., 2004) (Figure 4). As expected, failure loads generally increased with suture size for both loading conditions. We observed a significant effect of anchor design on *F*_*max*_ (suture size 0, p<0.001, F=108.825; suture size 2, p=0.001, F=15.394, suture size 5, p<0.001, F=14.861), but no significant effect of loading condition (p=0.862, F=0.031; p=0.373, F=0.841; p=0.212, F=1.644, for suture sizes 0, 2 and 5 respectively).

**Figure 4.**
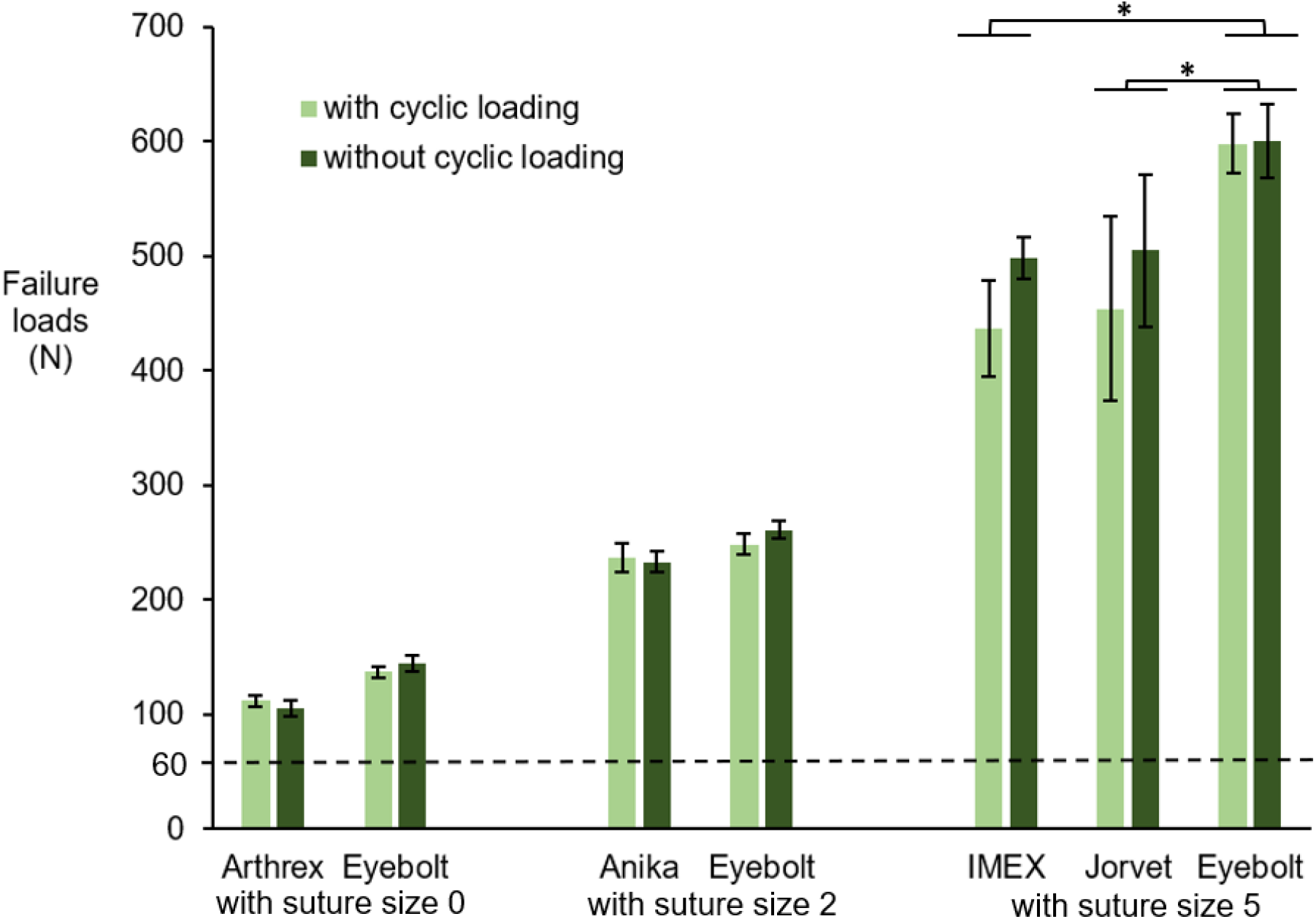
Failure loads of suture anchors compared with eyebolt screws, with accompanying suture sizes (n=5 per device and loading condition). Error bars represent standard deviations. Dashed line represents the reported peak Achilles tendon force in rabbit during hopping gait (West et al., 2004). The suture size used with each device is listed under the anchor name. *p<0.05

Compared to the eyebolt screw with corresponding suture sizes, *F*_*max*_ of the anchors were significantly less for the IMEX anchor (with cycling, p=0.002; without cycling, p=0.007) and Jorvet anchor (with cycling, p<0.001; without cycling, p=0.012).

All %Δ*F*_*max*_ values were negative, indicating that *F*_*max*_ for all suture anchors was always less than that of the eyebolt screw with the corresponding suture size (Figure 5). There was a significant effect of anchor design (p=0.003, F=5.623) but not loading condition (p=0.808, F=0.060) on %Δ*F*_*max*_. From post-hoc comparisons, %Δ*F*_*max*_ was significantly less for the Anika anchor than for the Arthrex (p=0.015), IMEX (p=0.004) and Jorvet (p<0.001) anchors.

**Figure 5.**
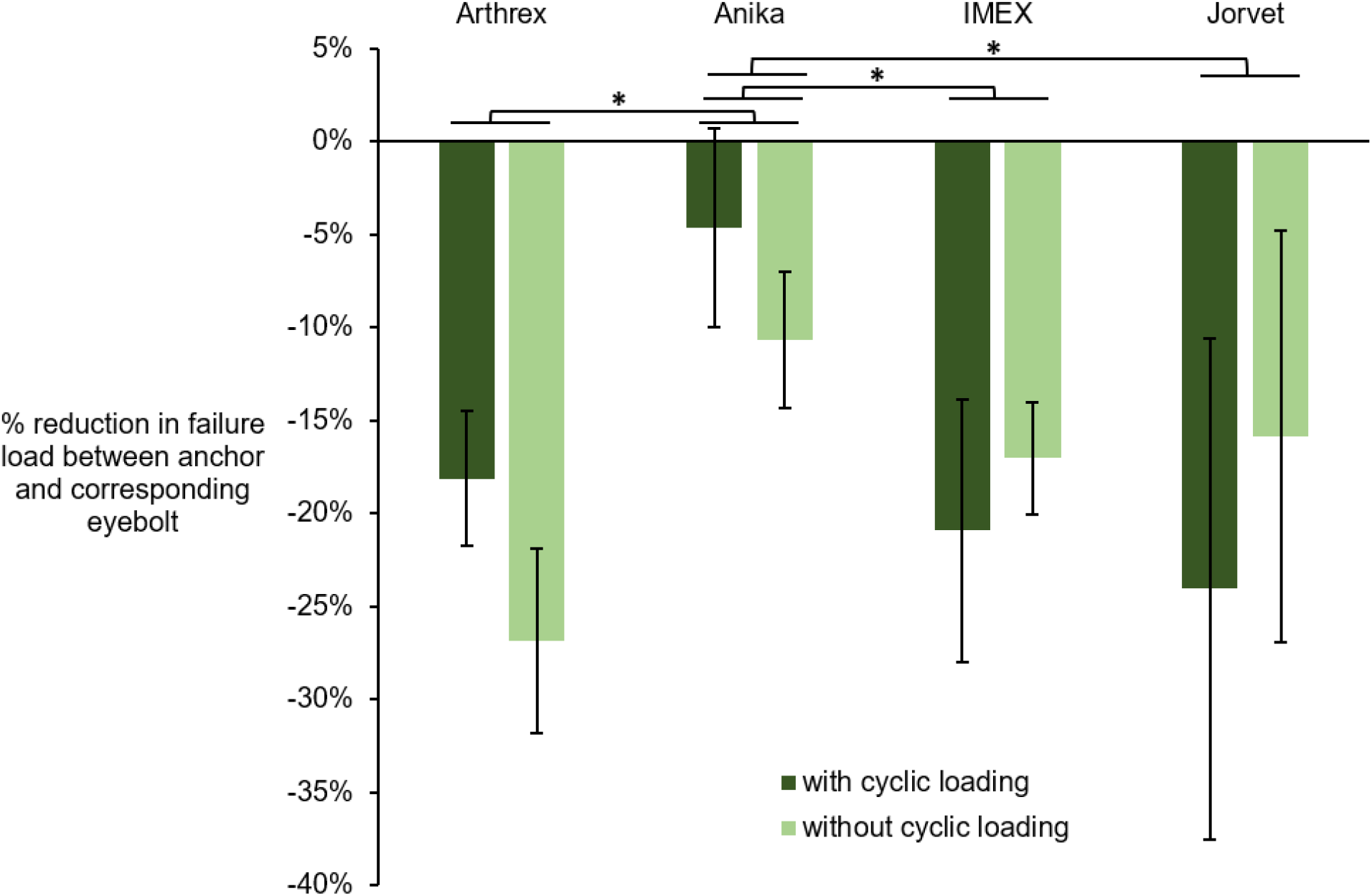
Percent difference in failure load between anchor and corresponding eyebolt using matched suture sizes (see Equation 1). Error bars represent standard deviations. *p<0.05

All suture anchors failed via suture breakage. For anchors with embedded eyelets (Anika and Arthrex) and the eyebolt screw, the suture broke at the knot in all samples (Figure 6). However, for the IMEX anchor, the suture failed at the mid-section under cyclic loading (n=2) and without cyclic loading (n=3); the remaining samples failed at the knot. For the Jorvet anchor, the suture broke at the mid-section for all five samples with cyclic loading and four samples without cyclic loading. The proportion of suture failure mode for the Jorvet anchor was significantly different from the corresponding eyebolt, both with cycling (p=0.024) and without cycling (p=0.004).

**Figure 6.**
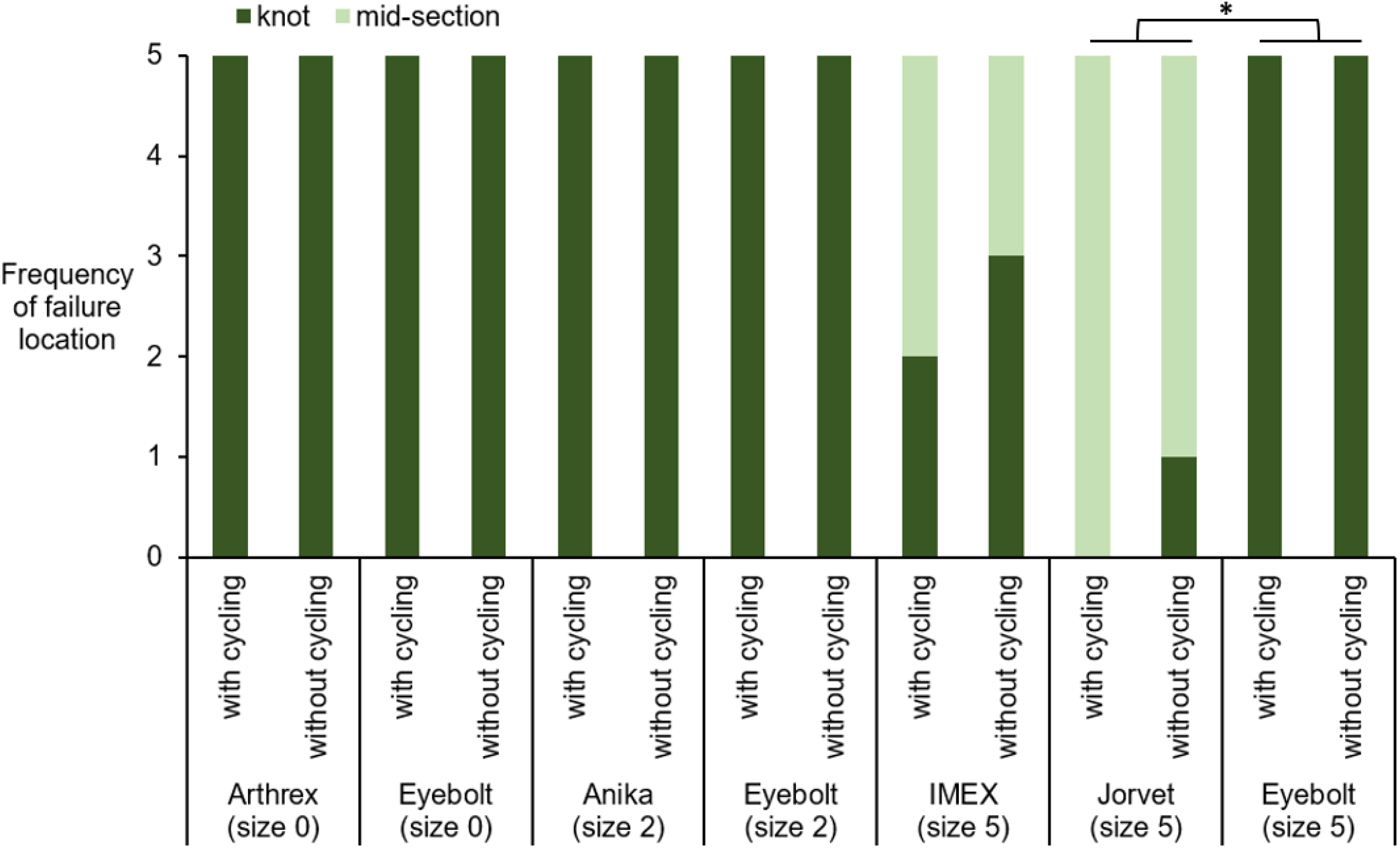
Frequency of suture failure mode (n=5 per device and loading condition). *p<0.05.

Conversely, the proportion of suture failure mode for the IMEX anchor was not significantly different from the corresponding eyebolt, both with cycling (p=0.083) and without cycling (p=0.222).

## DISCUSSION

The results generally supported our hypothesis 1 that the suture anchors with a raised eyelet would have a greater adverse effect on *F*_*max*_ compared to the suture anchors with an embedded eyelet. We identified from a previous study, the ultimate tensile strength of four anchors with raised eyelets and tested with stainless steel wire as the suture material (Barber et al., 1995) and compared these strengths of the suture anchor from the study with publicly available tensile strength of the stainless steel wire (Component supply, 2024). The reported ultimate strength in the study was observed to be less for three out of four anchors. This observation is consistent with the results of our study. In our study, *F*_*max*_ for all four suture anchors (Arthrex, Anika, IMEX and Jorvet) was significantly different from the respective eyebolt screw although loading conditions showed no significant effect. However, for %Δ*F*_*max*_, the Anika anchor had a significantly less adverse effect on *F*_*max*_ than the other three suture anchors we tested.

Across all suture anchors, cyclic loading up to 1000 cycles had no significant effect on *F*_*max*_ of the suture anchors for both anchors with raised eyelets and those with embedded eyelets.

However, in support of hypothesis 2, there was a stronger trend of an effect of cyclic loading for the anchors with raised eyelets (IMEX and Jorvet) than for anchors with embedded eyelets (Arthrex, Anika) (Figure 4). In a previous relevant study where the effect of cyclic loading was quantified (Balara et al., 2004), no effect of cyclic loading was observed after cycling metallic screw-type veterinary anchors for up to 1200 cycles. However, in a different study (Giles III et al., 2008), when suture anchors were tested for up 10,000 cycles, there were significant differences in the failure loads, suggesting that number of cycles to failures may be an important factor in determining the effect of cyclic loading on the strength of suture anchors.

Our results generally supported our hypothesis 3 that anchors with raised eyelets would have more frequent failures at the suture mid-section than anchors with embedded eyelets (Figure 6). When a suture anchor fails anywhere other than the knot, it raises concerns such as loading conditions and anchor design, because the knot is considered the weakest point in the construct and therefore the clinically expected point of failure. In a previous evaluation of suture anchors in human cadaveric shoulders (Barber et al., 1993), no significant difference in the failure loads of four suture anchors was reported. However, the mode of failure of the anchors varied from location to location. The predominant mode of failure in the proximal third of the humerus was via suture rupture whereas, in the greater tuberosity, one anchor failed via suture breakage and two other anchors failed via anchor pullout. This suggests that location of implantation may have an effect on suture anchor failure mode and that some suture anchors may be more suitable for some applications than others. Perhaps, this may be the case for the IMEX anchors used to implant artificial Achilles tendons in our rabbit model.

Taken together, our results suggest that the anchors with raised eyelets may have had design features that had a greater adverse effect on suture wear and strength than the anchors with embedded eyelets, depending on the application. For example, sutures placed in anchors with raised eyelets follow a less tortuous path than in anchors with embedded eyelets. Thus, sutures in raised eyelets may experience more relative motion, greater focus points of strain, and friction against edges, leading to more rapid suture wear. Given the more tortuous path, sutures in embedded eyelets may contact the anchor over a larger surface area and, thus, have greater load distribution (i.e., less load concentration). Finally, the tested anchors with raised eyelets may have had edges with smaller radii, which would also cause greater load concentration on the suture. Greater concentration of load on the suture is expected to result in greater (1) suture wear due to greater friction force and (2) probability of mid-section failure of the suture. To confirm these hypotheses, future studies should quantify the geometry of the suture anchors and relative motion between the suture and anchor during cyclic loading. Additionally, computational finite-element analysis may enable estimation of suture force magnitude, direction, and distribution for different anchors and loading conditions.

Our results only partly explain the suture anchor failures experienced in our previous in vivo study. The mechanical testing showed that IMEX suture anchors may be more prone to causing sutures to fail at the mid-section. However, the IMEX failure load exceeded those expected during in vivo peak Achilles tendon load by a factor of 8, and with no significant effect of cyclic loading. There are a few factors that may explain the discrepancy between the in vivo study and mechanical tests. For example, in our mechanical tests, the direction of the load was static. Conversely, in vivo, the suture anchors likely experienced more dynamic loading conditions, including simultaneous changes in magnitude and/or direction of tendon force relative to the suture anchor, due to dynamic ankle motion, muscle force, and dynamic external loads (i.e., ground reaction forces). Such dynamic loading may cause considerably more rubbing and wear of the suture against the anchor than in our mechanical tests. A related factor is the orientation of the suture anchor relative to the bone, which affects the direction of tendon load on the suture anchor; significant detrimental effects of anchor angulation and rotation on suture abrasion have been reported in a previous study (Bardana et al., 2003). In our vivo model, the anchors were typically inserted at an angle to the bone surface, though it is not clear if anchors rotated in the bone and the extent of such rotation.

An important consideration for selecting a suture anchor is the application or conditions for which it is designed, tested, and used. Several studies have characterized the ultimate strength or fatigue strength of suture anchors and in different applications including repair of anterior Bankart lesions, stabilization of anterior labral lesions, stabilization of canine hip joint luxation, rotator cuff repair and glenoid labral repair (Burkhart et al., 1997; Cummins & Murrell, 2003; Mueller et al., 2005; Nho et al., 2010; Rudzki et al., 2004). These tests were either with cyclic loading or without cycling. Failure in cyclic loading occurs due to material fatigue caused by repeated loading and unloading cycles and this test is used to determine the fatigue strength and fatigue life of a material whereas uniaxial load to failure is used to assess the acute mechanical strength of a material (McFarland et al., 2005). In human and animal models, suture anchors are mostly expected to be loaded in a dynamic manner. However, in most laboratory testing of these anchors, the loading conditions are static, with or without cycling. One study that involved dynamic loading did not measure strength of the anchor but rather evaluated two surgical techniques, with different number of anchors and anchor orientation (Garcia et al., 2013). The result suggests that the methods for evaluating the mechanical properties of suture anchors (static or dynamic loading) may depend on the specific application.

Publicly available product information for the anchors in this study provided limited information for applications or indications. For example, the company webpage for the IMEX anchor shows the device as a veterinary anchor with a representation of the anchor being used for the stabilization of cranial cruciate deficient stifles in what appears to be a small animal model. Detailed information regarding case outcomes is lacking and the use of these anchors may need to be more specifically studied relative to indications and applications best suited to their use (IMEX Veterinary Inc.). In our experience, use of these anchors may not be suitable for use in (re)attachment of the Achilles tendon to the calcaneus. A previous research indicated a connection between application and anchor strength (Barber et al., 1993). When immobilization following surgery is required and possible, the initial strength requirement of the suture anchorage may be small compared to those situations were immediate motion is required.

Our study had several limitations. First, the number of suture anchors and number of samples of each anchor included in our study were relatively small. Second, the depth of insertion of the anchor in the simulated bone block may have differed from the depth in vivo. Anchor depth has been implicated in the mechanical performance of suture anchor with deep sitting anchors reported to have higher failure loads and significantly different modes of failure compared with anchors with eyelets sitting above the bone (Bynum et al., 2005). Bone blocks are typically used for testing orthopedic implants, but their properties differ from those of rabbit bone. Several studies have tested suture anchors in human and animal cadaveric bone specimens, but we tested the anchors in vitro using bone blocks and not rabbit bones. Lastly, the conditioned and controlled laboratory conditions under which the mechanical testing was performed differ from the biological environment in which suture anchors are used clinically.

## CONCLUSION

The ultimate tensile strength of the anchors tested in this study exceeded the peak Achilles tendon force in rabbits during hopping gait and increased with increasing size of the accompanying suture. There are significant effects of anchor type but not loading condition, on the strength of the suture anchors. Of the anchors that were tested in this study, overall comparisons of suture anchors with embedded eyelets versus suture anchors with raised eyelets generally showed that raised eyelets had a greater adverse effect on (1) the maximum tensile force at failure, (2) the durability against cyclic loading, (3) the likelihood of failure at the suture mid-section, which is indicative of suture wear or cutting against the anchor. Future studies of suture anchors are needed to determine the influence of dynamic loading and its interaction with other factors such as eyelet position and geometry, on the strength and durability of the suture.

## Acknowledgement

this study was funded by the NIH (R61AR078096). We also wish to thank Elizabeth Croy and Dr Katrina Easton for their help during data collection.

